# What influences treatment response in animal models of non-alcoholic fatty liver disease? A meta-analysis with meta-regression

**DOI:** 10.1101/2019.12.31.887919

**Authors:** Harriet Hunter, Dana de Gracia Hahn, Amedine Duret, Yu Ri Im, Qinrong Cheah, Jiawen Dong, Madison Fairey, Clarissa Hjalmarsson, Alice Li, Hong Kai Lim, Lorcán McKeown, Claudia-Gabriela Mitrofan, Raunak Rao, Mrudula Utukuri, Ian A. Rowe, Jake P. Mann

## Abstract

The classical drug development pipeline necessitates studies using animal models of human disease to gauge future efficacy in humans, however, there is a comparatively low conversion rate from success in animals to in humans. Non-alcoholic fatty liver disease (NAFLD) is a complex chronic disease without any licensed therapies and hence a major field of animal research. We performed a meta-analysis of 414 interventional rodent studies (6,575 animals) in NAFLD to assess the mean difference in hepatic triglyceride content. 20 of 21 studied drug classes had similar efficacy with a mean difference of −30% hepatic triglyceride. However, when publication bias was accounted for, this reduced to −16% difference. Study characteristics were only able to account for a minority of variability on meta-regression, and we replicated previous findings of high risk of bias across 82% of cohorts. These findings build on previous work in preclinical neuroscience and help to explain the challenge of reproducibility and translation within the field of metabolism.

## Introduction

Interventional studies in animals are an integral component of the drug development pipeline. If a disease can be suitably modelled in an animal, then the therapeutic response to a treatment observed in animals should inform about its potential efficacy in humans[1]. However, there is a well-documented translational gap between preclinical studies and subsequent outcomes in humans[2–4]. Multiple factors contribute to this, including bias within study design[5], insufficiently powered preclinical studies[6], and biological differences between species[7,8].

Systematic analyses of preclinical studies, predominantly in the field of neuroscience, have found that publication bias may account for at least a third of the estimate of efficacy in trials[9,10]. In addition, other variables of animal model design can influence the magnitude of the treatment response[11]. These findings are highly relevant in the context of the ‘reproducibility crisis’[12,13] as well as having ethical implications of the use of animals in research that is not of optimum quality[14].

Non-alcoholic fatty liver disease (NAFLD) is a highly active field of animal research[15,16]. NAFLD is a common condition characterised by increased liver fat (hepatic steatosis) that may progress to inflammation (non-alcoholic steatohepatitis (NASH)) and fibrosis[17]. Cirrhosis, end-stage liver disease, and hepatocellular carcinoma develop in a small proportion of patients. However, due to the high prevalence of obesity, NAFLD is the second most common indication for liver transplant in the United States[18], predicted to overtake hepatitis C. NAFLD is intricately related with insulin resistance and therefore usually coexists with other features of the metabolic syndrome, such as type 2 diabetes and its recognised complications cerebrovascular disease, coronary artery disease and chronic kidney disease[19].

There are currently no approved pharmacological therapies for NAFLD[20]. Several Phase 3 trials are ongoing[21], but many interventions that appeared to have substantial efficacy in preclinical models have failed to be replicated in humans[22–24]. These studies have used a wide range of preclinical NAFLD models, including genetically modified animals (e.g. leptin deficient *ob/ob* mice), hypercaloric diets (e.g. high fat diet), and toxic insults (e.g. streptozocin injections), all of which may be used in varying combinations[25]. It is not known if, or which of, these variables influence treatment response to therapeutic agents in preclinical models of NAFLD.

Therefore, we performed a meta-analysis of interventional rodent studies of NAFLD to describe which drug classes were associated with the greatest reduction in liver fat and whether any study characteristics (or biases) were linked to the magnitude of effect.

## Results

We performed a systematic search to identify interventional studies in rodent models of NAFLD. Our searches yielded 7503 articles, which after screening gave 4467 articles for full-text review (Figure 1). Studies were included in the meta-analysis if they used a pharmacological agent that had been used in Phase 2 or 3 trials in humans for NAFLD and reported hepatic triglyceride content for control and interventional groups. 244 studies were included in the meta-analysis, comprising 414 cohorts of rodents (6,575 animals).

**Fig. 1:**
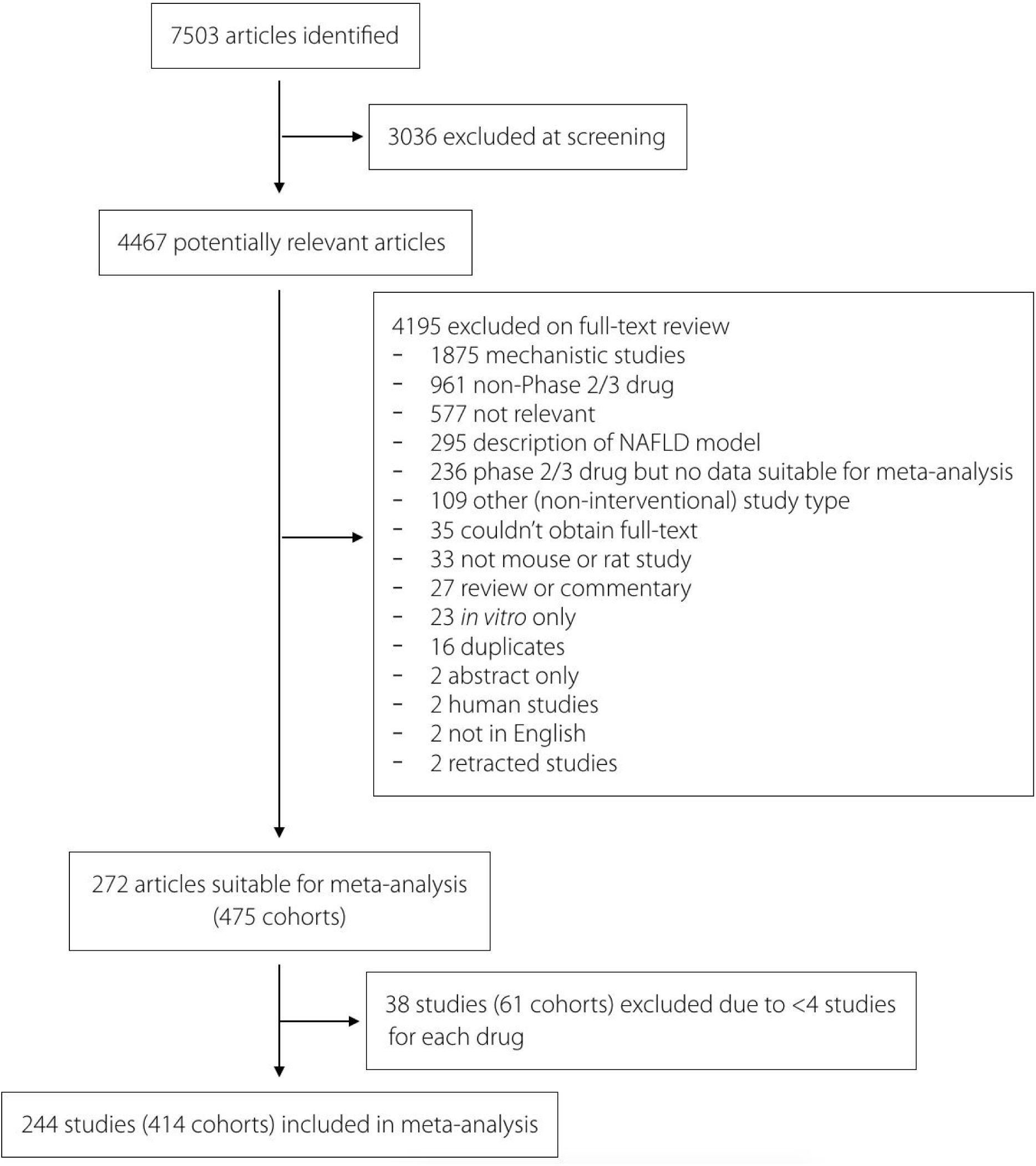
Study inclusion and exclusion flow chart.

21 drug classes were represented in the meta-analysis, including 34 studies on GLP-1 agonists, 15 on probiotics, and 55 on fibrates.

### Meta-analysis of hepatic triglyceride content

We used random-effects meta-analysis to estimate the mean difference in hepatic triglyceride (TG) content between intervention and control groups. The overall mean change in hepatic TG content was −30.4% (95% CI −33.0%, −27.7%) with considerable between-study heterogeneity (I^2^ = 91% [95% CI 90.3-91.6%]).

We hypothesised that much of this heterogeneity would be due to the different drug class interventions. However, on meta-analysis using drug class as a sub-group, we found marked similarity between the effect size of the different interventions (Figure 2). There was weak evidence of difference between drug classes (Q=36.4, p=.014). The confidence intervals of 20 out of 21 drug classes overlapped and there remained substantial or considerable heterogeneity within drug class subgroups (Table 1).

**Table 1:**
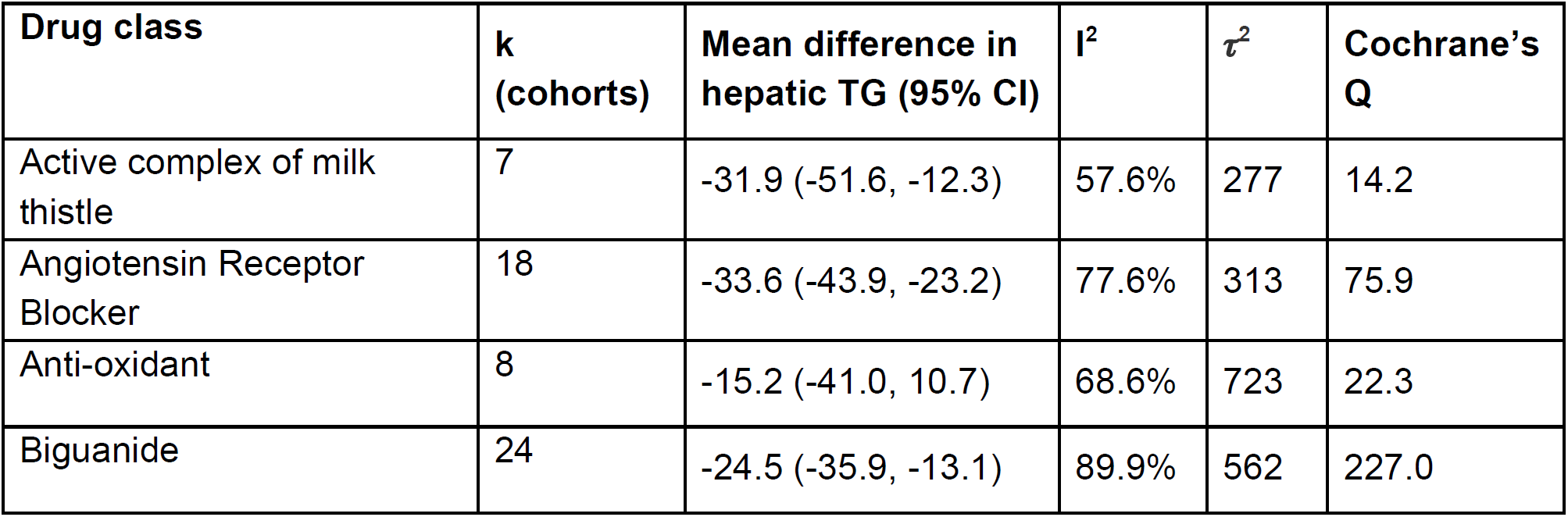
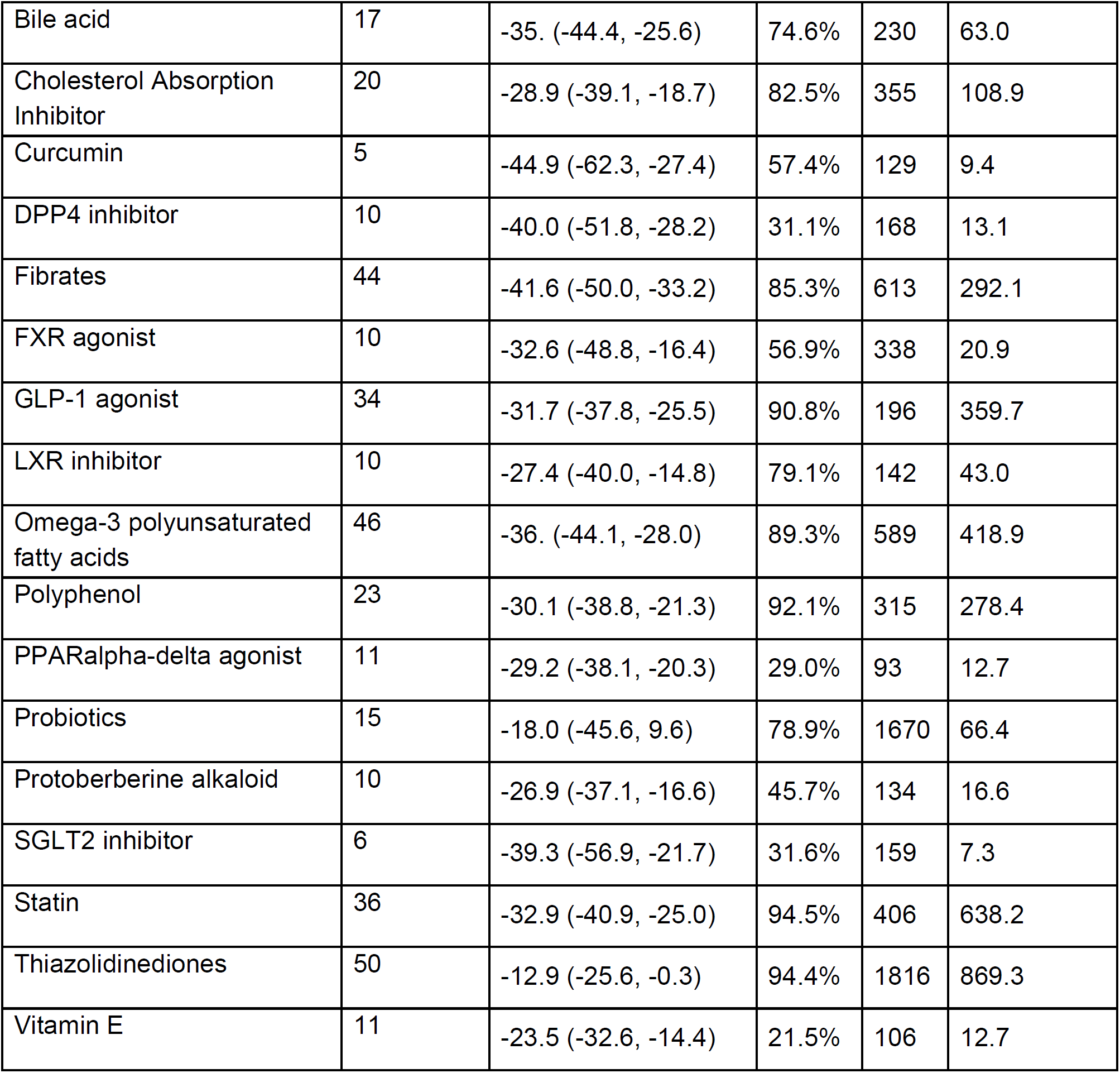
Summary of the studies by drug class with estimates of within sub-group heterogeneity. DPP4, dipeptidyl peptidase-4; FXR, farnesoid X receptor; GLP-1, glucagon-like peptide-1; LXR, liver X receptor; PPAR, peroxisome proliferator-activated receptor; SGLT2, sodium-glucose transport protein 2.

| Drug class | k (cohorts) | Mean difference in hepatic TG (95% CI) | $I^2$ | $\tau^2$ | Cochrane's Q |
| --- | --- | --- | --- | --- | --- |
| Active complex of milk thistle | 7 | -31.9 (-51.6, -12.3) | 57.6% | 277 | 14.2 |
| Angiotensin Receptor Blocker | 18 | -33.6 (-43.9, -23.2) | 77.6% | 313 | 75.9 |
| Anti-oxidant | 8 | -15.2 (-41.0, 10.7) | 68.6% | 723 | 22.3 |
| Biguanide | 24 | -24.5 (-35.9, -13.1) | 89.9% | 562 | 227.0 |
| Bile acid | 17 | -35. (-44.4, -25.6) | 74.6% | 230 | 63.0 |
| Cholesterol Absorption Inhibitor | 20 | -28.9 (-39.1, -18.7) | 82.5% | 355 | 108.9 |
| Curcumin | 5 | -44.9 (-62.3, -27.4) | 57.4% | 129 | 9.4 |
| DPP4 inhibitor | 10 | -40.0 (-51.8, -28.2) | 31.1% | 168 | 13.1 |
| Fibrates | 44 | -41.6 (-50.0, -33.2) | 85.3% | 613 | 292.1 |
| FXR agonist | 10 | -32.6 (-48.8, -16.4) | 56.9% | 338 | 20.9 |
| GLP-1 agonist | 34 | -31.7 (-37.8, -25.5) | 90.8% | 196 | 359.7 |
| LXR inhibitor | 10 | -27.4 (-40.0, -14.8) | 79.1% | 142 | 43.0 |
| Omega-3 polyunsaturated fatty acids | 46 | -36. (-44.1, -28.0) | 89.3% | 589 | 418.9 |
| Polyphenol | 23 | -30.1 (-38.8, -21.3) | 92.1% | 315 | 278.4 |
| PPARalpha-delta agonist | 11 | -29.2 (-38.1, -20.3) | 29.0% | 93 | 12.7 |
| Probiotics | 15 | -18.0 (-45.6, 9.6) | 78.9% | 1670 | 66.4 |
| Protoberberine alkaloid | 10 | -26.9 (-37.1, -16.6) | 45.7% | 134 | 16.6 |
| SGLT2 inhibitor | 6 | -39.3 (-56.9, -21.7) | 31.6% | 159 | 7.3 |
| Statin | 36 | -32.9 (-40.9, -25.0) | 94.5% | 406 | 638.2 |
| Thiazolidinediones | 50 | -12.9 (-25.6, -0.3) | 94.4% | 1816 | 869.3 |
| Vitamin E | 11 | -23.5 (-32.6, -14.4) | 21.5% | 106 | 12.7 |

**Fig 2.**
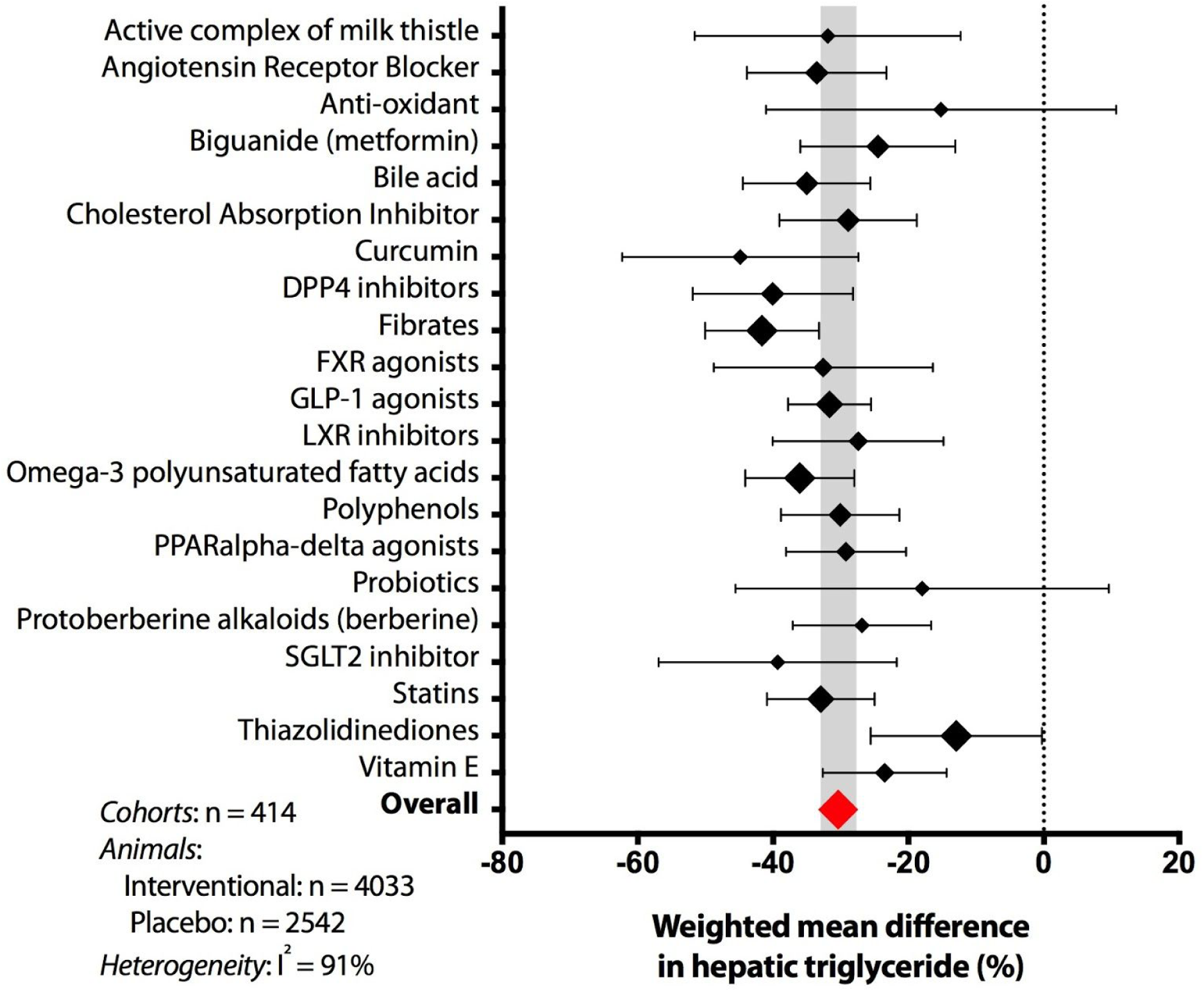
Summary forest plot showing mean difference in hepatic triglyceride content between interventional and control animals in rodent studies of NAFLD. Individual studies have been hidden and only subgroup summary figures are illustrated.

In order to test whether the heterogeneity was driven by individual outlying studies, we used a Baujat plot to identify outliers with disproportionate contribution to heterogeneity (Supplementary Figure 1). After removal of seven studies, the overall mean difference was unchanged (−31% [95% CI −33%, −28%]) and considerable heterogeneity remained (I^2^ = 86.5% [95% CI 85.3%, 87.5%]).

### Meta-regression for difference in hepatic TG content

To try and understand the variability in mean difference of hepatic TG, we performed meta-regression using both categorical and continuous variables associated. On mixed-effects meta-regression, drug class accounted for only 5.6% of heterogeneity in hepatic TG (Table 2).

**Table 2.** Results from univariable meta-regressions. β represents the change in mean difference in hepatic TG per unit change in variable. For categorical variables, the effect sizes are shown in Figure 2 (for drug class), Supplementary Table 1 (genetic background), and Supplementary Table 2 (model type).

| Variable(s) | k (Number of cohorts) | $\beta$ (SE) | Test of moderators (p-value) | $r^2$ |
| --- | --- | --- | --- | --- |
| Drug class | 414 | Figure 2 | 0.002 | 5.60% |
| Genetic background | 337 | SupTable 1 | 0.08 | 1.65% |
| Age at end of study (months) | 392 | 1.45 (.60) | 0.02 | 1.13% |
| Duration of intervention (weeks) | 413 | .42 (.22) | 0.06 | 0.63% |
| Age at start of drug (months) | 392 | .94 (.73) | 0.2 | 0.12% |
| Model | 277 | SupTable 2 | 0.37 | 0.11% |
| Quality score [0-4] | 414 | 1.3 (1.8) | 0.46 | 0 |
| Male sex | 382 | -1.9 (26.6) | 0.68 | 0 |
| Female sex | 382 | 3.3 (27.2) | 0.68 | 0 |
| % fat in diet | 219 | -.06 (.15) | 0.66 | 0 |
| Drug dose (mg/kg) | 283 | .005 (.007) | 0.5 | 0 |

When other study characteristics and variables were assessed, genetic background (Supplementary Table 1) and age at the end of intervention (Figure 3) had a very modest impact upon mean difference in hepatic TG. The type of NAFLD model used accounted for almost no heterogeneity in the data (Supplementary Table 2).

**Figure 3.**
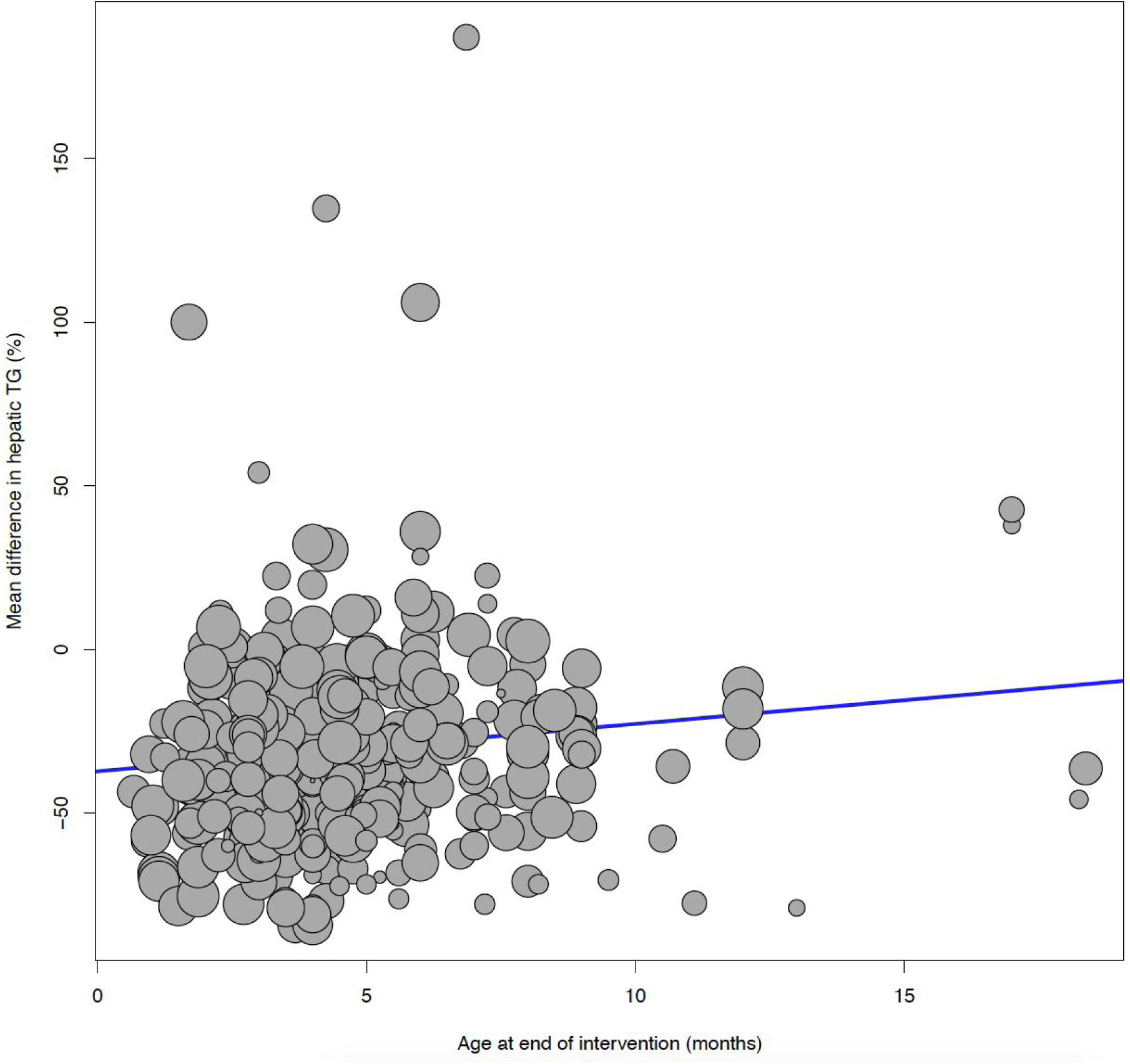
Bubble plot illustrating the results of meta-regression between the age at end of intervention (in months) and mean difference in hepatic TG. Studies in older mice were weakly associated with smaller treatment responses: 1.4% less reduction in hepatic TG per month of life for the studied rodents.

In order to estimate the total variance explained by measurable variables, we used multiple variable meta-regression in a smaller dataset (k = 71) with minimum 4 studies for each drug, genetic background, and NAFLD model. When all variables from Table 2 were included, this model was not associated with treatment response (F25,45 = .07, p = .83) though accounted for 47.4% of heterogeneity.

### Publication (study distribution) bias

The meta-analysis result was assessed for study bias using a funnel plot (Figure 4). This showed an uneven distribution with a bias towards positive results (i.e. a reduction in hepatic TG with intervention), which was supported by Egger’s test (β = −1.0 [95% CI −1.4, −.60], p = 6×10^−5^). Using the Trim and Fill method to account for this bias, we estimated that the true overall mean difference in hepatic TG would be half as great: −16.6% (95% CI −19.6, −13.7).

**Fig 3.**
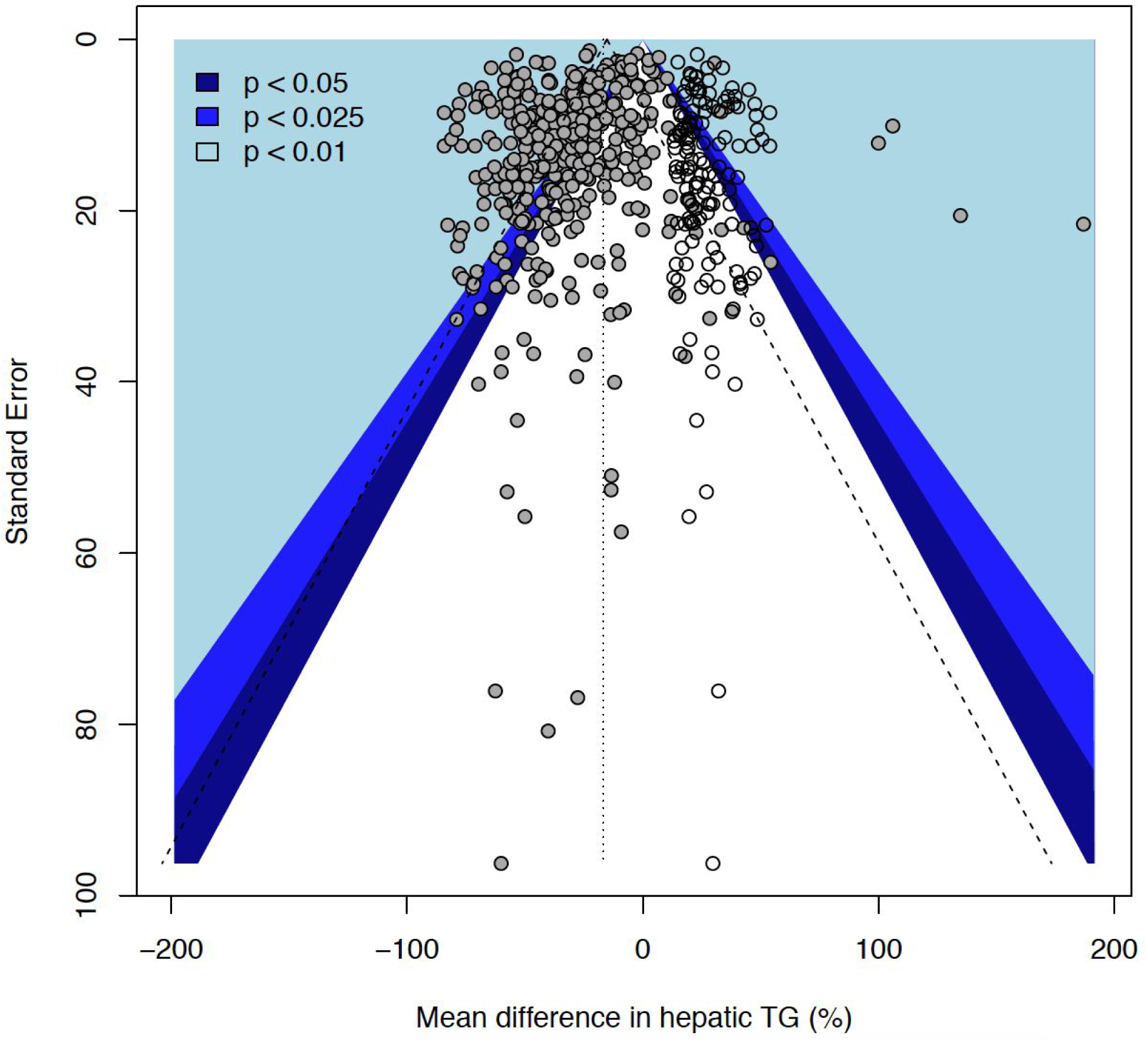
Funnel plot illustrating study distribution (publication) bias in 414 original studies (solid grey circles) with 139 added studies (from trim and fill). The statistical significance associated with each study is illustrated with the coloured background.

### Study quality

We used a four-item scale to estimate study quality (Supplementary Figure 2). We found that 361/441 (82%) cohorts were at high risk of bias due to either absence of randomisation or absence of blinding. No individual component of the risk of bias score (or combined as a study quality score) was associated with magnitude of treatment response.

There was a bimodal distribution of study powers across the cohorts included in the meta-analysis (Supplementary Figure 3). 91% (375/414) cohorts had a power of 80% or greater on post-hoc calculation. However, using the results from this meta-analysis (mean difference −30.4% with standard deviation = 29.6), to achieve a power of 80% with significance set as p=.05, group size would need to be n=16. 3.4% (14/414) cohorts included 16 or more animals and would have met sufficient power to detect associations, based on these data.

## Discussion

Through meta-analysis and meta-regression we have quantified the remarkable similarity between interventional drug classes and which study characteristics influence treatment response in rodent models of NAFLD. These findings suggest that measurable factors (e.g. animal age) can only account for a small degree of variability in animal response to drugs.

NAFLD is generally considered to be a major public health concern and drug development is a highly active field[26]. Though there are no licensed therapies, there have been over 30 drugs used in Phase 2 or 3 trials, with some demonstrating potential efficacy in well-conducted randomized controlled trials, such as for GLP-1 agonists[27] and pioglitazone[28]. Therefore, we were surprised to find that there was minimal difference between any of 21 drug classes studied in animal models (Figure 2). In fact, thiazolidinediones were the only outlier, due to studies that found an increase in hepatic TG content in response to treatment[29,30]. To our knowledge this is such similarity between multiple drugs in preclinical models has not been described before to this extent. It is generally consistent with findings reported in preclinical models of spinal cord injury where the effect size of several different types of treatment overlapped[11].

Despite overlapping confidence intervals for drug classes, there was considerable heterogeneity between studies. This is potentially expected as a consequence of including a relatively large number of cohorts in the meta-analysis (n=414). We were further surprised to find that drug class accounted for a modest amount of variability in treatment response. Similarly, it is well established that different rodent NAFLD models result in higher or lower hepatic TG concentrations[25], but our results do not suggest that they influence treatment response. Selection of animal models for studying NAFLD is a heavily debated topic[31] and a detailed discussion is beyond the scope of this meta-analysis.

Design of an interventional animal study is considerably more complex than just the ‘core’ model (e.g. leptin deficient (*ob/ob*) mice). There have been previous reports to illustrate the effect of genetic background on NAFLD[32,33] as well as in other fields, including immunology[34] and behavioural neuroscience[35,36]. We were surprised to find that genetic background accounts for only 1.6% heterogeneity in results. We anticipate that the true value is greater than this because we could only include backgrounds that had been used in multiple studies and we excluded mixed genetic backgrounds from analysis.

Older animals had a smaller response to treatment, which is consistent with evidence that they develop more substantial NASH[37] and their physiology is less plastic[38]. However, it could be argued that younger animals do not accurately reflect chronic multisystem diseases such as NAFLD.

Using the trim and fill method, we estimated that study distribution bias (most likely publication bias in this case) may have almost doubled the reported magnitude of effect (16% reduction in hepatic TG compared to 30%). The presence of publication bias did not come as a surprise[39] and this dataset provides useful replication of the strong evidence base for this in preclinical neurological studies. The results from power calculations are also likely to reflect publication bias: based on the overall effect summary, only 3.4% of cohorts were of sufficient size to be predicted to achieve the power of 80%. However, over 90% of cohorts had results that would be consistent with power >80%. Similarly, we have replicated previously described low rates of randomization and blinding in animal studies[40].

There are several implications of these results. Firstly, it is not surprising that there are multiple reports of difficulty in reproducing preclinical studies of metabolism[13] if the choice of drug intervention plays a relatively minor role in determining the magnitude of treatment response. Variations in study design affects response and therefore could silence subtle differences, especially when combined with this risk of bias which can generate false positives. Secondly, these results also help to explain the difficulty in bridging the preclinical to human translational gap[41]. If almost all studied interventions are reported to have a similar effect in animals, then results to date are not helpful in guiding which will show efficacy in humans. This is clearly a challenge across the entire biomedical field and careful replication of animal study design is likely to be important. It could be suggested that 30% reduction in hepatic TG could be a ‘benchmark’ for future therapeutics aimed at reduced steatosis.

This apparent threshold for 30% reduction in hepatic TG does not appear to be due to the sensitivity of in vitro assays used as TG content was reported to range from 0.05µM/mg to 7.9µM/mg across the included studies.

Much of the variability in results could not be accounted for. We speculate that unmeasurable variables, such as technique of animal handling, may also influence the treatment effect. The bacterial status of mice is known to affect liver phenotypes[42], potentially via intestinal dysbiosis[43,44]; and similarly, whilst we included proportion of fat in diet as a variable, the complete diet composition is much more complex and may be of importance.

The main strength of this work is the number of included studies, interventions, and variables. This has facilitated a detailed analysis of a single disease area. Though the primary limitation of findings is that we are unable to determine the extent that these results are generalisable to other fields we have highlighted several conceptual similarities to previous meta-research in neuroscience. In addition, many studies did not report variables, for example genetic background of animals was reported in 7% (31/441), which reduced the number of studies included in meta-regression analyses.

## Conclusion

Quantifiable and measurable variables in animal studies, including the drug used, have only a modest effect on the size of treatment response. There is a highly consistent 30% reduction in hepatic TG in all drugs tested to date, though the true magnitude of this value might be half as great when accounting for publication bias. Standardisation of study design and rigorous quality are needed in preclinical studies in metabolism to improve the translation and replicability of findings.

## Methods

### Review protocol and search strategy

The systematic review protocol was prospectively registered with SyRF (Systematic Review Facility) and is available from: https://drive.google.com/file/d/0B7Z0eAxKc8ApQ0p4OG5SblRlRTA/view.

PubMed via MEDLINE and EMBASE were searched for published articles of experimental rodent models of fatty liver, NAFLD, or non-alcoholic steatohepatitis (NASH). The following search term was used: (“Non-alcoholic fatty liver disease” OR “Nonalcoholic fatty liver disease” OR “NAFLD” OR “non-alcoholic steatohepatitis” OR “nonalcoholic steatohepatitis” OR “NASH” OR “fatty liver” OR “hepatic steatosis”) AND (“mouse” OR “animal” OR “rat” OR “murine” OR “animal model” OR “murine model” OR “rodent model” OR “experimental model”) NOT (“Review”). Both databases were searched using the “Animal” filters[45,46], the results combined, and duplicates eliminated. The search was completed in November 2017.

### Study selection and eligibility criteria

Our inclusion criteria were as follows: primary research articles using mice or rats to model NAFLD (to include hepatic steatosis, NASH, and NASH-fibrosis), use of pharmacological intervention with a control (or placebo) group, and that the pharmacological intervention class (e.g. statins) had been used in Phase 2 or 3 trials in humans for treatment of NAFLD/NASH. Studies were excluded if: not modelling NAFLD/NASH; studies in humans or any animal other than mice and rats; reviews, comments, letters, editorials, meta-analyses, ideas; articles not in English (unless there was an available translation); studies not reporting hepatic triglyceride content relative to hepatic protein (e.g. mg/mg or µM/mg); studies using a pharmacological agent class that had not been used in Phase 2/3 studies in humans for NAFLD; and fewer than four independent studies using any single pharmacological agent drug class.

Abstracts and titles were screened to identify relevant studies using Rayyan[47]. Potentially relevant studies had their full text extracted and were assessed against inclusion/exclusion criteria independently by two reviewers, with discrepancies settled by discussion with JPM.

### Data collection

The variables extracted were as follows: phenotypic characteristics of animal model used (sex, diet [including percentage of fat in diet], rodent age, genetic alterations, background animal strain); drug treatment (dose, drug class, duration, age at intervention), and hepatic triglyceride content. Studies frequently included multiple cohorts or interventional arms, which were defined as use of a different animal model of NAFLD, a different drug, or a different drug dose. Data were extracted for each cohort or interventional arm separately.

### Quality assessment

Each paper was assessed in the following 4 areas: use of a protocol, reporting use of randomisation, reporting use of blinding, and a power calculation. These were each given a score of 1, and each paper was assigned an overall “quality score”. A post-hoc power calculation was performed for each study using the means of each group and a common SD [48] using the *pwr[49]* package in r. In addition, a ‘pre-test’ sample size calculation was performed using the overall effect summary from meta-analysis, power = 80% and p-value = .05.

### Statistical analysis

Primary outcome was the percentage difference in hepatic triglyceride (TG) content in the interventional group compared to control/placebo.

Random-effects meta-analysis using the Hartung-Knapp-Sidik-Jonkman method was used to calculate mean difference in hepatic TG for each drug class. Heterogeneity within drug classes and across the whole dataset was reported using Cochran’s Q, Higgin’s & Thompson’s I^2^, and *τ*^2^. Potential outliers were identified using a Baujat plot[50], where all studies with excess contribution to heterogeneity were excluded as a sensitivity analysis.

Study distribution (‘publication’) bias was assessed using funnel plot with Egger’s test. Given evidence of study distribution bias, Duval & Tweedie’s trim-and-fill procedure[51] was performed to estimate the impact of bias on the overall measure.

Mixed-effects meta-regression was performed to assess which baseline variables were associated with heterogeneity in TG content across the whole dataset. Meta-regression was performed for categorical variables (drug class, sex, animal background, NAFLD model design) and continuous variables (percentage of fat in diet, age at intervention, drug dose). For each regression analysis, studies were only included where four or more studies reported each variable. For example, for analysis by animal background, C57BL/6J (used in 110 studies) was included but FVB/N (used in 2 studies) was excluded. Due to high variability and minimal replication, studies using ‘Mixed’ animal background were excluded from meta-regression analyses. The number of cohorts included in each regression analysis is reported with their results. Multiple variable meta-regression was performed to assess what proportion of between-study heterogeneity could be accounted for by baseline characteristics (using R^2^).

Statistical analysis was performed using R 3.6.2 for Mac[52,53] with packages *dmetar[53], meta[54]*, and *metafor[55]*. Graphs were also generated using GraphPad Prism (v8.0 for Mac, GraphPad Software, La Jolla California, USA).

## Data availability

The raw dataset used for analysis, including references to individual studies, are available in the Dryad repository: https://doi.org/10.5061/dryad.pzgmsbcgc

R code used for analysis are available in Supplementary Data.

