## Supplementary data for "What influences treatment response in animal models of non-alcoholic fatty liver disease? A meta-analysis with meta-regression"

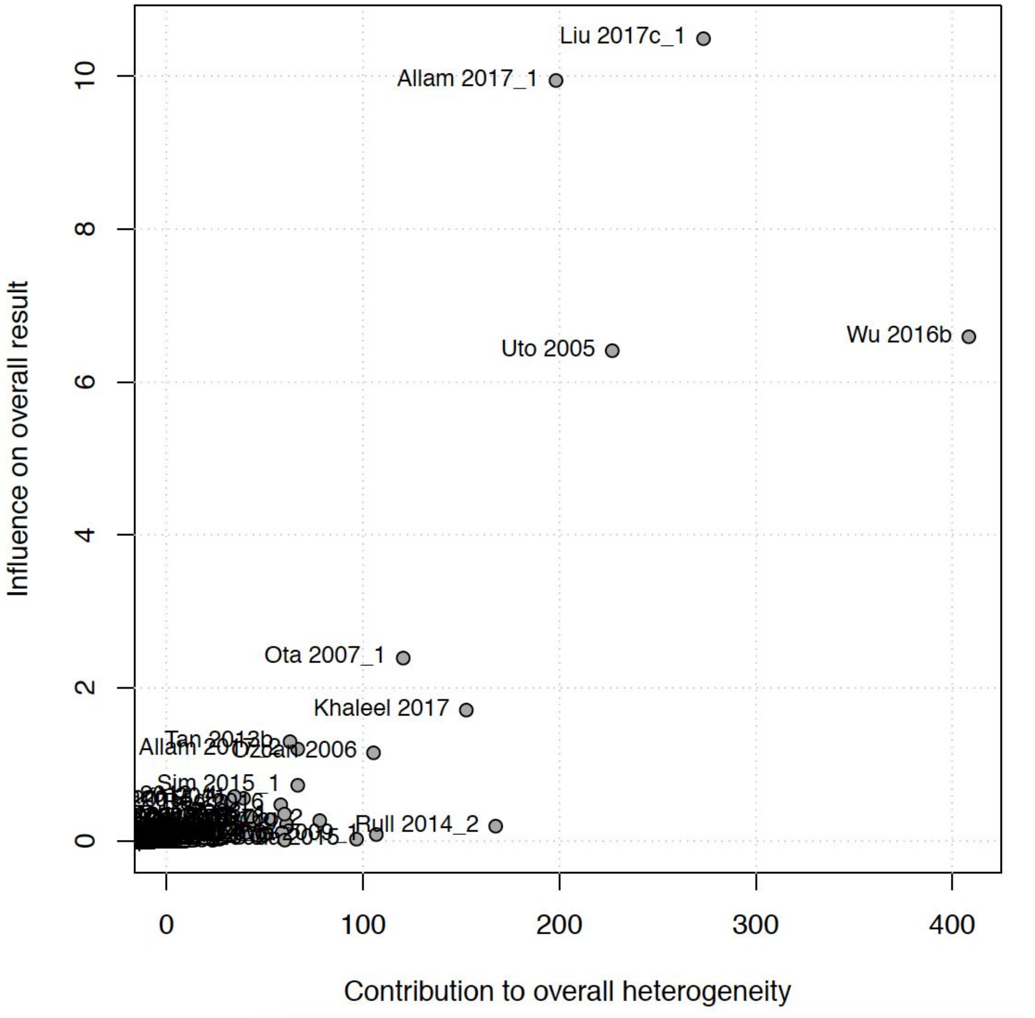


***Supplementary Figure 1.*** *Baujat plot showing individual study contributions to heterogeneity in the meta-analysis. The studies with highest contribution were excluded as a sensitivity analysis.*

| Genetic background | *β* (SE) | p-value |
|---|---|---|

| | | |
|---|---|---|
| C57BL/6J | 3.4 (4.5) | 0.45 |
| C57BLKS/J | -9.1 (11.3) | 0.42 |
| ICR | -18.2 (8.3) | 0.03 |
| KK-Ay | 15. (9.6) | 0.12 |
| Long Evans | -2.6 (10.1) | 0.80 |
| Sprague Dawley | -4. (5.4) | 0.46 |
| Wistar | 0.6 (4.9) | 0.91 |
| Zucker | 12.7 (8.9) | 0.15 |

***Supplementary Table 1.*** *Summary of results from meta-regression using animal genetic background as a categorical variable. β represents the change in mean difference in hepatic TG associated with each model type. SE, standard error.*

| **Model** | ***β* (SE)** | **p-value** |
|---|---|---|
| Fructose | -13.7 (9.7) | 0.16 |
| HFD + STZ | -19. (13.7) | 0.17 |
| High Fat + High Cholesterol Diet | -5.3 (9.7) | 0.59 |
| High Fat + High Fructose Diet | 3.0 (11.0) | 0.79 |
| High Fat + High Sucrose | -7.9 (9.8) | 0.42 |
| High Fat Diet / HFD | -7.8 (8.3) | 0.35 |
| Leptin Deficiency (ob/ob) | -16.2 (11.8) | 0.17 |
| Methionine and choline deficient diet / MCD | -8.6 (8.7) | 0.33 |
| Otsuka Long-Evans Tokushima Fatty (OLETF) rat | -24.3 (15.1) | 0.11 |

| | | |
|---|---|---|
| Zucker | 4.7 (10.8) | 0.66 |

***Supplementary Table 2.*** *Summary of results from meta-regression using NAFLD model as a categorical variable. β represents the change in mean difference in hepatic TG associated with each model type. SE, standard error.*

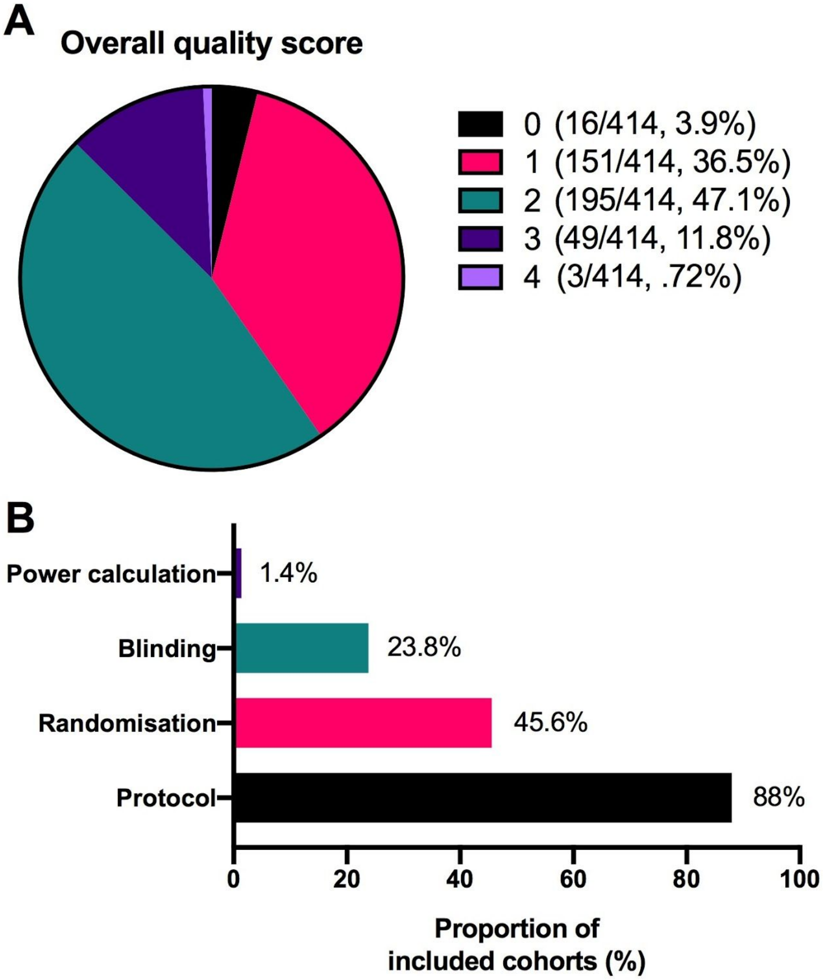


***Supplementary Figure 2.*** *Quality score used to describe the risk of bias of included cohorts. A four-point scale was composed of the use of a power calculation, use of blinding, randomisation, and referring to a predefined protocol, with 1-point awarded for presence of each factor. Overall quality score from 0-4 is shown in A and proportion of cohorts achieving each factor is shown in B.*

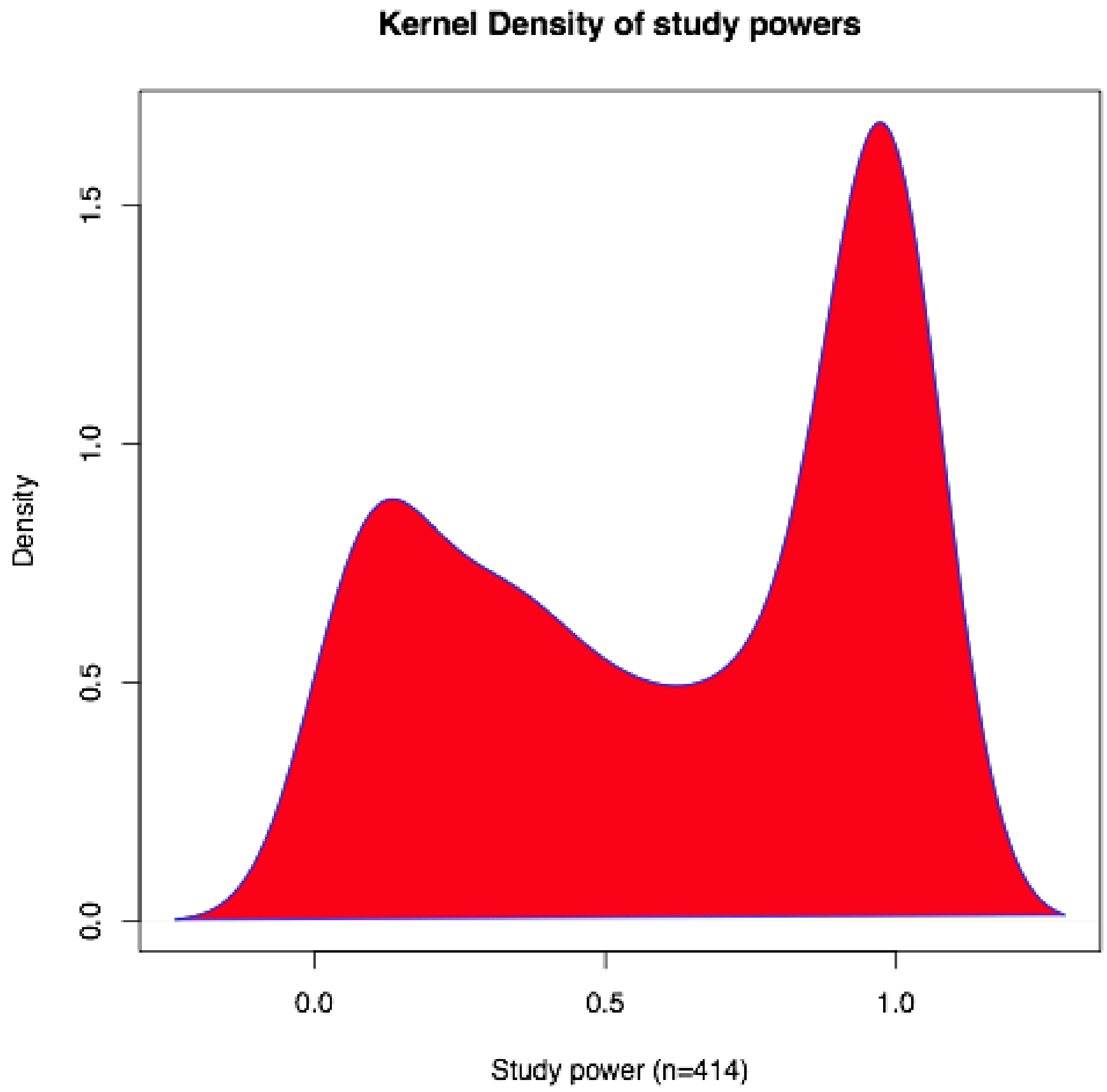


***Supplementary Figure 3.*** *Kernel density plot illustrating the distribution of post-hoc calculated study powers for cohorts included in the meta-analysis.*

### Supplementary methods

**R code used in analysis**

```
install.packages("tidyverse")
install.packages("meta")
install.packages("metafor")
install.packages("devtools")
devtools::install_github("MathiasHarrer/dmetar")
library(dmetar)
```

```
library(meta)
library(metafor)
library(readxl)
max.print = 100000

## perform meta-analysis of hepatic TG with sub-group analysis by drug class
TG_drug <- read_excel("TG_drug.xlsx")
m.tg.drug <- metacont(TG_Int_size, TG_Int_Mean, TG_Int_SD, TG_Pla_size, TG_Pla_Mean,
TG_Pla_SD, data = TG_drug, studlab = paste(Paper), comb.fixed = FALSE, comb.random =
TRUE, method.tau = "SJ", hakn = TRUE, prediction = TRUE, sm = "MD")
drug.subgroup <- update(m.tg.drug, byvar = Drug_Class, bylab = "Drug Class")
pdf(file="drug_subgroup_v1.pdf",width=15,height=120)
forest(drug.subgroup)
dev.off()
sink("m_tg_drug.txt")
print(drug.subgroup)
sink()

## identify outliers
pdf(file="baujat_drug_v1.pdf")
baujat(m.tg.drug)
dev.off()

## use dataset with 7 outliers removed and repeat meta-analysis
TG_outlier <- read_excel("TG_outlier.xlsx")
m.tg.outlier <- metacont(TG_Int_size, TG_Int_Mean, TG_Int_SD, TG_Pla_size, TG_Pla_Mean,
TG_Pla_SD, data = TG_outlier, studlab = paste(Paper), comb.fixed = FALSE, comb.random =
TRUE, method.tau = "SJ", hakn = TRUE, prediction = TRUE, sm = "MD")
m.tg.outlier
drug.outlier <- update(m.tg.outlier, byvar = Drug_Class, bylab = "Drug Class")
drug.outlier
sink("m_tg_outlier.txt")
print(drug.outlier)
sink()

## perform univariable meta-regressions
reg.drug <- metareg(m.tg.drug,Drug_Class)
reg.drug
sink("reg_drug.txt")
print(reg.drug)
sink()

reg.fat <- metareg(m.tg.drug,Fat_kcal)
```

```
reg.fat
sink("reg_fat.txt")
print(reg.fat)
sink()

reg.age_start <- metareg(m.tg.drug,Age_start)
reg.age_start
sink("reg_age_start.txt")
print(reg.age_start)
sink()

reg.age_end <- metareg(m.tg.drug,Age_end)
reg.age_end
sink("reg_age_end.txt")
print(reg.age_end)
sink()
pdf(file="bubble_age_end.pdf",width=10,height=10)
bubble(reg.age_end, xlab = "Age at end of intervention (months)", ylab = "Mean difference in
hepatic TG (%)", col.line = "blue", lwd = 3)
dev.off()

reg.duration <- metareg(m.tg.drug,Duration)
reg.duration
sink("reg_duration.txt")
print(reg.duration)
sink()

reg.sex <- metareg(m.tg.drug,Sex)
reg.sex
sink("reg_sex.txt")
print(reg.sex)
sink()

reg.qual <- metareg(m.tg.drug,Qual_score)
reg.qual
sink("reg_qual.txt")
print(reg.qual)
sink()


## use dataset where there is at least 4 studies for each genetic background
TG_bg <- read_excel("TG_bg.xlsx")
```

```
m.tg.bg <- metacont(TG_Int_size, TG_Int_Mean, TG_Int_SD, TG_Pla_size, TG_Pla_Mean,
TG_Pla_SD, data = TG_bg, studlab = paste(Paper), comb.fixed = FALSE, comb.random =
TRUE, method.tau = "SJ", hakn = TRUE, prediction = TRUE, sm = "MD")
bg.subgroup <- update(m.tg.bg, byvar = Background_simple, bylab = "Background")
sink("m_tg_bg.txt")
print(bg.subgroup)
sink()
reg.bg <- metareg(m.tg.bg,Background_simple)
reg.bg
sink("reg_bg.txt")
print(reg.bg)
sink()

## use dataset where there is at least 4 studies for each NAFLD model type
TG_model <- read_excel("TG_model.xlsx")
m.tg.model <- metacont(TG_Int_size, TG_Int_Mean, TG_Int_SD, TG_Pla_size, TG_Pla_Mean,
TG_Pla_SD, data = TG_model, studlab = paste(Paper), comb.fixed = FALSE, comb.random =
TRUE, method.tau = "SJ", hakn = TRUE, prediction = TRUE, sm = "MD")
model.subgroup <- update(m.tg.model, byvar = Models, bylab = "Model")
sink("m_tg_model.txt")
print(model.subgroup)
sink()
reg.model <- metareg(m.tg.model,Models)
reg.model
sink("reg_model.txt")
print(reg.model)
sink()

## use dataset containing drug dose data
TG_dose <- read_excel("TG_dose.xlsx")
m.tg.dose <- metacont(TG_Int_size, TG_Int_Mean, TG_Int_SD, TG_Pla_size, TG_Pla_Mean,
TG_Pla_SD, data = TG_dose, studlab = paste(Paper), comb.fixed = FALSE, comb.random =
TRUE, method.tau = "SJ", hakn = TRUE, prediction = TRUE, sm = "MD")
reg.dose <- metareg(m.tg.dose,Drug_dose)
reg.dose
sink("reg_dose.txt")
print(reg.dose)
sink()

## multiple variable meta-regression
## extract effect summaries from original meta-analysis
m.tg.drug_output <- data.frame(m.tg.drug$TE,m.tg.drug$seTE,m.tg.drug$studlab)
write.table(m.tg.drug_output,file="drug_output.csv",sep=",")
```

```
## combine yi and sei from m.tg.drug with meta-data from TG_drug to generate TG_mmetareg
and filter studies for only those with >4 of each drug, model, and background
TG_mmetareg2 <- read_excel("TG_mmetareg2.xlsx")
TG_mmetareg2$Models = factor(TG_mmetareg2$Models)
TG_mmetareg2$Intervention = factor(TG_mmetareg2$Intervention)
TG_mmetareg2$Drug_Class = factor(TG_mmetareg2$Drug_Class)
TG_mmetareg2$Background_simple = factor(TG_mmetareg2$Background_simple)
rma_all3 <- rma(yi = yi, sei = sei, data = TG_mmetareg2, method = "ML", mods = ~ Models +
Age_start + Age_end + Drug_Class + Background_simple + Drug_dose + Duration + Sex +
Qual_score + Fat_kcal, test="knha")
rma_all3
sink("mreg_all.txt")
print(rma_all3)
sink()

## perform bias analysis
egg.drug <- eggers.test(x = m.tg.drug)
egg.drug
sink("eggers_drug.txt")
print(egg.drug)
sink()
m.tg.trimfill <- trimfill(m.tg.drug)
m.tg.trimfill
sink("trim_fill.txt")
print(m.tg.trimfill)
sink()
pdf(file="trimfill_funnel_v6.pdf")
funnel(m.tg.trimfill, xlab="Mean difference in hepatic TG (%)", contour = c(.95,.975,.99),
col.contour=c("darkblue","blue","lightblue")) + legend(-200, 1, c("p < 0.05", "p < 0.025", "p <
0.01"), bty = "n", fill=c("darkblue","blue","lightblue"))
dev.off()


##Power analysis

install.packages("pwr")
library(pwr)
install.packages("square")
library(square)

##calculate common SD
TG_drug$Int_SD2 <- TG_drug$TG_Int_SD*TG_drug$TG_Int_SD
```

```
TG_drug$Pla_SD2 <- TG_drug$TG_Pla_SD*TG_drug$TG_Pla_SD
TG_drug$comm_SD <- sqrt(((TG_drug$Int_SD2+TG_drug$Pla_SD2)/2))
TG_drug$m_diff <- TG_drug$TG_Pla_Mean-TG_drug$TG_Int_Mean
TG_drug$eff_size <- TG_drug$m_diff/TG_drug$comm_SD


TG_drug$power <- pwr.t2n.test(n1 = TG_drug$TG_Int_size, n2 = TG_drug$TG_Pla_size, d =
TG_drug$eff_size)$power

write.table(TG_drug,file="TG_drug_power.csv",sep=",")

hist(TG_drug$power)

dens_power <- density(TG_drug_power$power)

pdf(file="kernel_power_v1.pdf")
plot(dens_power, main="Kernel Density of study powers", xlab = "Study power (n=414)")
polygon(dens_power, col="red", border="blue")
dev.off()

##calculate sample size needed for average results
##mean comm_SD = 29.587
##meta-analysis mean difference = -30.4
##cohen's d = -30.4/29.587 = -1.02746

pwr.t.test(d = -1.02746, power = 0.80, sig.level = 0.05)
```
